# Histone deacetylase 5 in prelimbic prefrontal cortex limits context-associated cocaine seeking

**DOI:** 10.1101/2024.09.21.614125

**Authors:** Sarah M. Barry, Jessica Huebschman, Derek M Devries, Lauren M McCue, Evgeny Tsvetkov, Ethan M. Anderson, Benjamin M. Siemsen, Stefano Berto, Michael D. Scofield, Makoto Taniguchi, Rachel D. Penrod, Christopher W. Cowan

**Author notes:** Correspondence, Medical University of South Carolina Department of Neuroscience, 173 Ashley Avenue, BSB 403, MSC 510, Charleston, SC 29425, USA.

## Abstract

**Background:** Repeated cocaine use produces neuroadaptations that support drug craving and relapse in substance use disorders (SUDs). Powerful associations formed with drug-use environments can promote a return to active drug use in SUD patients, but the molecular mechanisms that control the formation of these prepotent drug-context associations remain unclear.

**Methods:** In the rat intravenous cocaine self-administration (SA) model, we examined the role and regulation of histone deacetylase 5 (HDAC5) in the prelimbic (PrL) and infralimbic (IL) cortices in context-associated drug seeking. To this end, we employed viral molecular tools, chemogenetics, RNA-sequencing, electrophysiology, and immunohistochemistry.

**Results:** In the PrL, reduction of endogenous HDAC5 augmented context-associated, but not cue-or drug prime-reinstated cocaine seeking, whereas overexpression of HDAC5 in PrL, but not IL, reduced context-associated cocaine seeking, but it had no effects on sucrose seeking. In contrast, PrL HDAC5 overexpression following acquisition of cocaine SA had no effects on future cocaine seeking. We found that HDAC5 and cocaine SA altered the expression of numerous PrL genes, including many synapse-associated genes. HDAC5 significantly increased inhibitory synaptic transmission onto PrL deep-layer pyramidal neurons, and it reduced the induction of FOS-positive neurons in the cocaine SA environment.

**Conclusions:** Our findings reveal an essential and selective role for PrL HDAC5 to limit associations formed in cocaine, but not sucrose, SA environments, and that it alters the PrL excitatory/inhibitory balance, possibly through epigenetic regulation of synaptic genes. These results further position HDAC5 as a key factor regulating reward-circuit neuroadaptations that underlie common relapse triggers in SUD.

## Introduction

Substance use disorders (SUDs) are a significant public health challenge and are associated with a reduced quality of life and increased mortality. Cocaine is the most abused stimulant-class drug, and deaths due to cocaine use have more than doubled since 2012^1^. Despite these challenges, no pharmacological agents have been approved for safe and effective treatment of cocaine use disorder (CUD)^2^. Repeated drug use causes transcriptional alterations in brain reward areas that support persistent, functional alterations that predispose individuals to drug-seeking and relapse behavior^3,4^. The medial prefrontal cortex (mPFC) is a critical component of drug-seeking circuitry due to its important role in executive functions, including learning, outcome predictions, memory retrieval, decision making, emotional processing, and cognitive control^5–10^. The mPFC can be divided into subdivisions based on their functions and projections along a dorsal to ventral axis^11–13^. Two subregions of the mPFC, the prelimbic cortex (PrL) and the infralimbic cortex (IL), undergo extensive alterations following use of illicit substances and are critical components of the circuitry involved in a return to active drug use (i.e., relapse)^14–19^. The mPFC also serves as a major glutamatergic input to the nucleus accumbens (NAc) and this projection is critically involved in drug-seeking behavior ^20–22^.

Neuroadaptations within the mPFC, including epigenetic alterations, contribute to persistent drug-seeking^23–25^. Post-translational modifications to histone tails are one way that gene expression is regulated following drug experiences. Histone deacetylases (HDACs) have been shown to mediate cocaine-induced changes in cellular functions^26–30^. Class IIa HDACs are unique in the HDAC family because synaptic activity, or activation of cAMP signaling, can trigger a shift in their nucleocytoplasmic localization^28^. One class IIa HDAC, HDAC5, is highly expressed in the medial prefrontal cortex (mPFC)^31,32^ and cocaine use causes a decrease of HDAC5 mRNA in the prefrontal cortex^33^. In addition, HDAC5 in the NAc serves to limit both cocaine conditioned place preference and cue- and drug prime-reinstated cocaine or heroin seeking following intravenous drug self-administration (SA)^28–30^. Indeed, non-contingent or contingent drug exposure induces a transient nuclear accumulation of HDAC5 in NAc MSNs, which appears to be driven in part by activation of dopamine D1 receptors, cAMP signaling, and dephosphorylation of three conserved HDAC5 serines^28^. Indeed, expression of a dephosphorylated, nuclear-localized HDAC5 mutant in the NAc (S259A/S279A/S498A or “HDAC5-3SA”) reduces the development of cocaine CPP and cued- and drug-primed cocaine-seeking following SA training^29^.

Despite strong cortical expression, HDAC5’s regulation and function in the mPFC, a key brain hub for drug-seeking behavior, has not been explored in SUD-related behavior. We show here that HDAC5 in PrL, but not IL, plays a selective role to limit context-associated cocaine, but not sucrose, seeking, and it regulates numerous PrL synaptic genes, GABAergic synaptic transmission, and FOS-induction in the cocaine SA environment.

## Results

### HDAC5 subcellular localization is regulated by synaptic activity in cortical neurons

In cultured primary cortical neurons, transfected wild-type HDAC5 was observed in both the cytoplasm and nucleus, but with a larger proportion of neurons showing a greater distribution within the cytoplasm. Since drugs of abuse activate dopamine signaling in multiple addiction-related brain regions, we tested whether cortical HDAC5 can be regulated by cAMP signaling. Similar to NAc neurons^28^, increasing cAMP signaling with forskolin (FSK) caused a translocation of HDAC5 to the nucleus in >80% of the neurons (**Fig 1A**, 2-way ANOVA, Significant Interaction, Treatment x Localization; F(1,4)= 113.3, *p=* -.0004***, Sidak’s Multiple Comparisons (MC), DMSO: *p=*0.0007, FSK: *p*= <0.0001).). We next infused a neurotropic AAV2-3XFlag-HDAC5 virus in the adult rat mPFC and assessed HDAC5 nucleocytoplasmic distribution following intravenous saline or cocaine self-administration (SA). On the seventh and final day of SA, the rats were sacrificed at 1 hour into their SA session (t=1 hr) or 2 hours after the end of a 2-hour session (t=4 hrs). Compared to saline SA, we observed that cocaine SA produced a significant, decrease in HDAC5’s cytoplasmic/nuclear distribution in mPFC layer V, but not layer II/III, neurons at the 4-hr, but not 1-hr, timepoint (**Fig 1**). Furthermore, there was a significant reduction in Layer V cytoplasmic localization of HDAC5, compared to Layers II/III, at 4-hrs following cocaine SA (**Fig 1B**, Ordinary one-way ANOVA, F(7, 268) = 4.664, *p<*0001; Planned Sidak’s Multiple Comparisons, Sal 4hr L-V vs Coc 4hr L-V, p=0.0021, Coc 4hr L-II/III vs Coc 4hr L-V, *p=*0.0053), suggesting that cocaine SA promotes partial nuclear accumulation of HDAC5 in deep-layer mPFC neurons.

**Figure 1.**
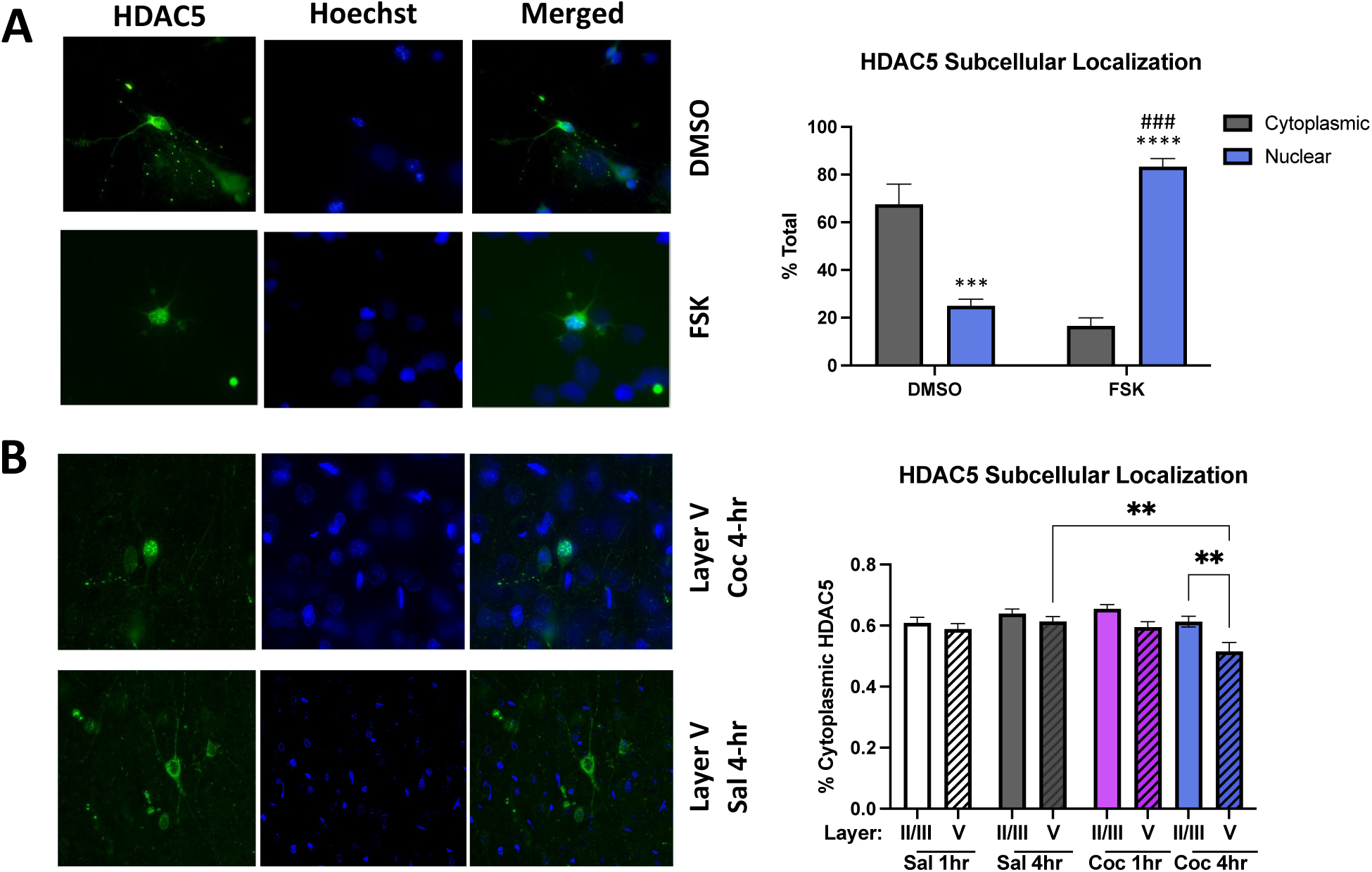
cAMP activation or cocaine SA promote PrL HDAC5 nuclear accumulation. (**A)** In cultured cortical cells, HDAC5 is primarily located in the cytoplasm and nucleus under control conditions. After a 3-hour incubation of 10mm Forskolin (FSK), HDAC5 is translocated to the nucleus. (**B)** *In vivo* expression of HDAC5-wt-Flag in the PrL of rats self-administering *i.v.* saline or cocaine, HDAC5 translocated to the nucleus in Layer V of cocaine self-administering rats at the 4-hr timepoint.

### HDAC5 in the prelimbic, but not infralimbic, prefrontal cortex limits context-associated cocaine seeking

To investigate HDAC5’s influence in mPFC on cocaine SA behavior, we infused in the adult PrL a re-validated AAV2-shHDAC5^30^ to reduce endogenous *Hdac5* mRNA in this brain region. Rats were then allowed to SA cocaine on an increasing fixed-ratio (FR) schedule (**Fig. 2A**). Compared to the shRNA negative control (AAV2-shLucif), viral-mediated HDAC5 knockdown in PrL had no effects on any aspect of cocaine SA acquisition, operant discrimination between the active vs. inactive lever, cocaine infusions, or stable drug-intake levels (**Figs. 2B**, Mixed-Effect Analysis, Day x Group, F(12, 107)= 0.625 (active lever), Mixed-Effect Analysis, Day x Group F(12, 107)= 0.9060 (inactive lever); **S1A,** Mixed-Effect Analysis, Day x Group F(12, 107)= 0.7648 (cocaine infusions)). However, following 7 days of home-cage abstinence, shHDAC5^PrL^ rats displayed a significant increase in cocaine seeking under non-reinforced extinction conditions (**Fig. 2C**, t(9)=2.58, *p=*0.323), which we refer to as “context-associated cocaine seeking”. Following extinction training, where the rats reduced their pressing of the drug-paired lever (**Fig. S1B,** Main Effect, Group F(1.=,9)= 8.593; Main Effect Day F(1.401, 12.61)= 47.30), shHDAC5^PrL^ rats displayed cue-reinstated cocaine seeking and cocaine-primed drug seeking that was indistinguishable from control rats (**Figs. 2D**, t(9)=0.165, *p=*0.8724; **S1C** t(9)=0.2467, *p=*0.817), suggesting that endogenous HDAC5 in the PrL limits context-associated cocaine seeking, but not cued or drug-primed reinstatement of cocaine seeking.

**Figure 2.**
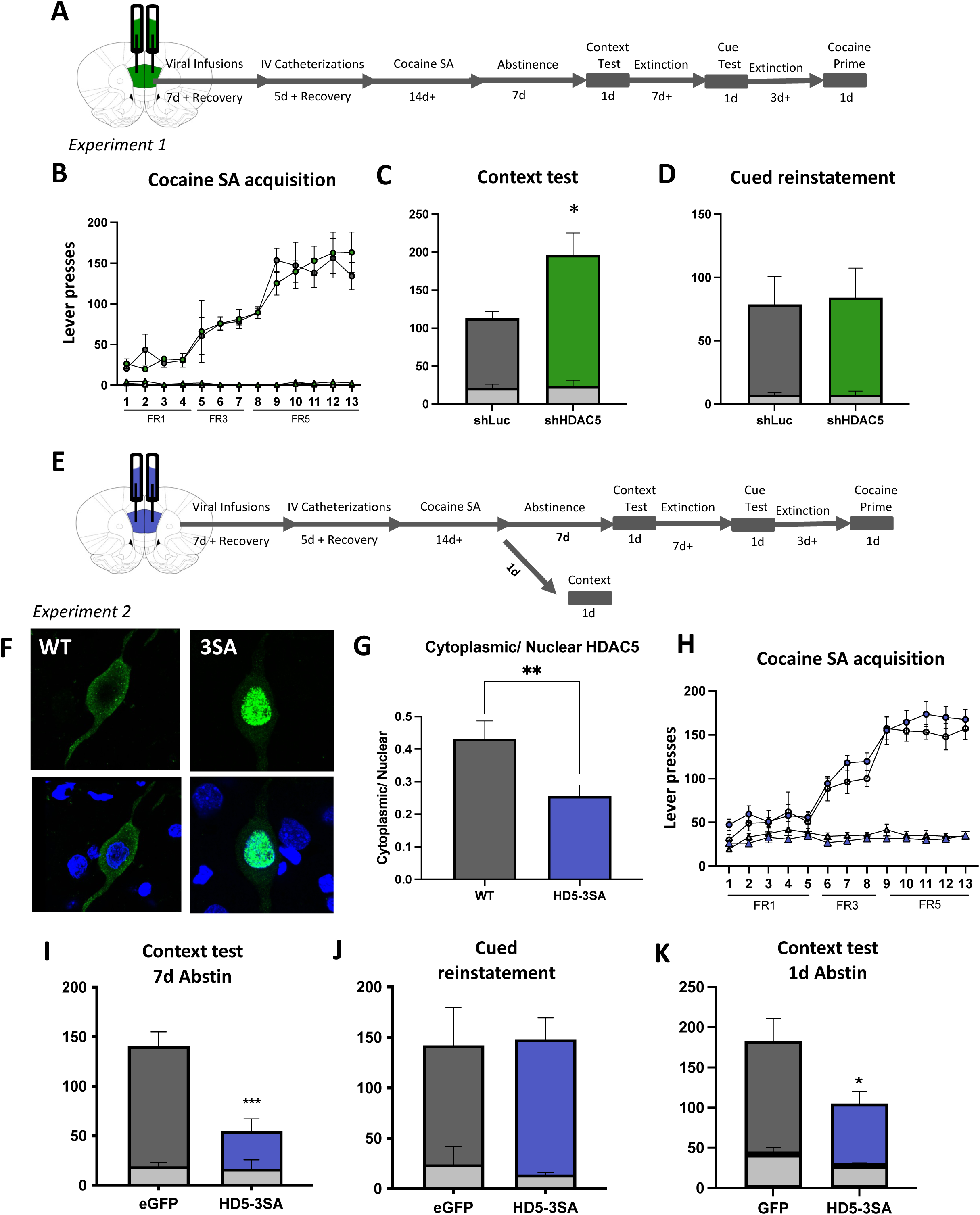
HDAC5 expression in the PrL bidirectionally regulates context associated drug-seeking in a drug-specific manner. (**A**) Experiment #1 timeline. (**B**) AAV-shHDAC5-mediated reduction of endogenous HDAC5 expression in the PrL did not alter cocaine SA acquisition or stable cocaine-taking behavior. (**C**) PrL shHDAC5 augmented context associated cocaine-seeking behavior compared to AAV2-shLuc control condition. (**D**) Following extinction training, there were no group differences in a cued reinstatement test. (**E**). Experiment #2 timeline with representative image of PrL HDAC5-3SA expression. (**F, G**) HDAC5-3SA shows increased nuclear localization in comparison to wild-type HDAC5. (**H**) HDAC5-3SA expression did not alter acquisition behaviors. (**I**) Following 7 days of abstinence, PrL HDAC5-3SA significantly suppressed context-associated cocaine seeking. (**J**) HDAC5-3SA expression had no effect on cue-reinstated seeking after extinction. (**K**) A subset of animals from Experiment #2 underwent the context test after a 24-hr abstinence, and HDAC5-3SA significantly reduced cocaine seeking in the operant context.

We next asked whether increasing nuclear HDAC5 levels in PrL could suppress context-associated cocaine seeking. To this end, we infused AAV2-3xFlag-HDAC5-3SA^29^, a nuclear-localized mutant of HDAC5, or AAV2-EGFP bilaterally in the PrL of adult rats and allowed them to SA cocaine (**Fig. 2 E-F,** HDAC5-3SA validation of cortical *in vivo* nuclear localization **Fig 2G** t(25)=2.876, *p=*0.0093). We found that there were no group differences in active or inactive lever pressing for cocaine (**Fig. 2H**) or in total cocaine taking (**Fig. S1D**). However, following 7 days of home-cage abstinence, HDAC5-3SA produced a significant reduction in context-associated cocaine seeking (**Fig. 2I,** t(11)=4.828, *p=*0.0005). Similar to shHDAC5, we detected no effects of PrL HDAC5-3SA overexpression on cued or drug-primed reinstatement of cocaine seeking (**Fig. 2J and Fig. S1E-F**), suggesting again a selective effect of PrL HDAC5 to limit context-related drug seeking. To test whether PrL HDAC5-3SA reduces context-associated seeking by limiting incubation of cocaine craving^34,35^ produced by the 1-week of home-cage abstinence, we tested rats in the context-associated seeking test after only 1 day of abstinence. Surprisingly, HDAC5-3SA still suppressed context-associated cocaine seeking after only 1 day of withdrawal (**Fig. 2K**, t(13)=2.549, *p*=0.0242), suggesting that extended withdrawal is not required for HDAC5 to limit drug-context associations. To test whether PrL HDAC5 limits general reward-related seeking, we next tested the effects of PrL HDAC5-3SA on sucrose SA and subsequent sucrose-seeking behavior. However, in contrast to cocaine SA, we detected no effects of PrL HDAC5 on context-associated sucrose seeking or any other aspect of sucrose SA behavior (**Figs. 3A-C** (sucrose SA: no significant main effect of group, F(10, 90)= 18.16, *p=*0.3747) **and S2A-B**), suggesting that HDAC5-3SA’s effects on context-associated seeking are drug-specific.

**Figure 3.**
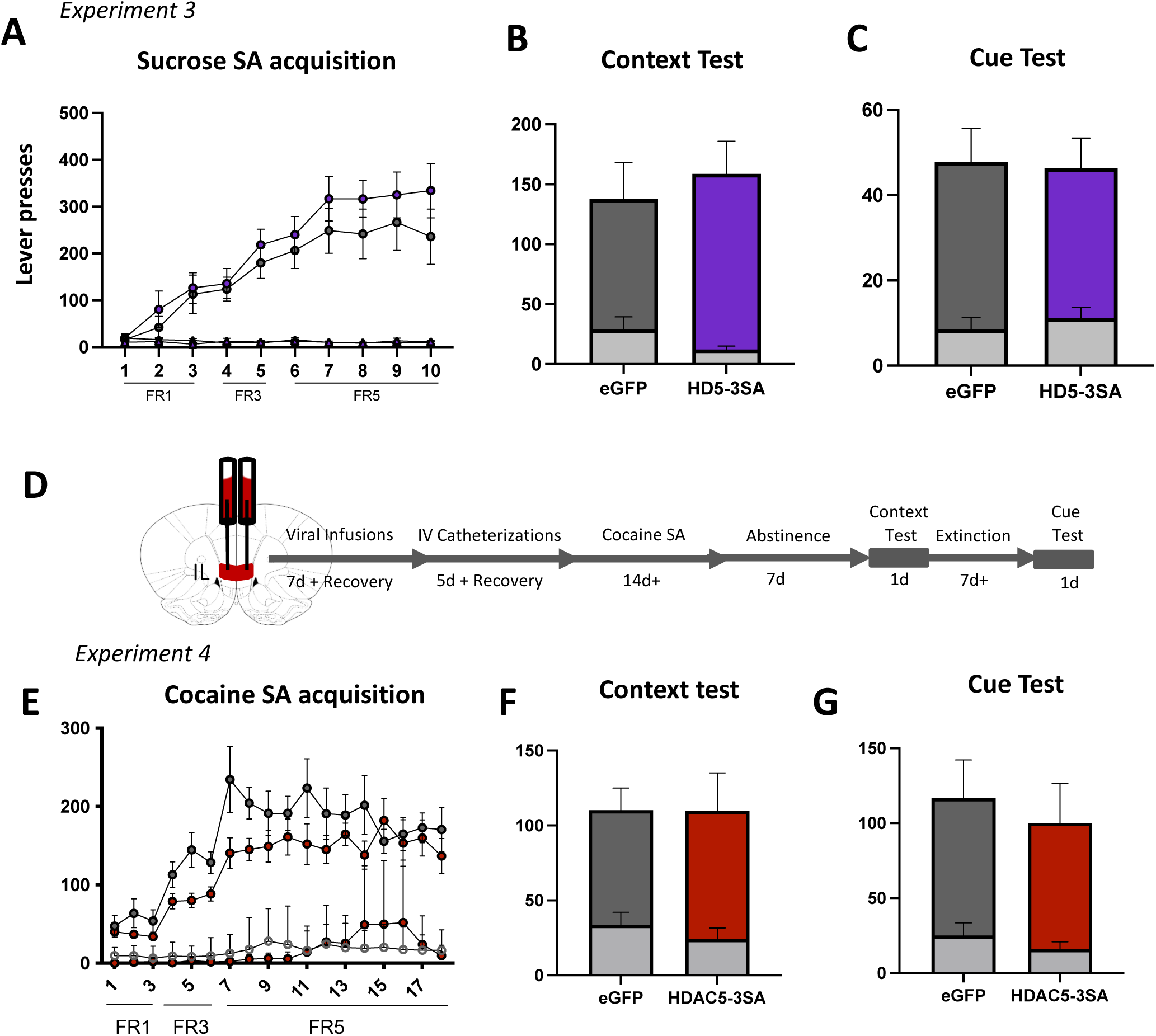
HDAC5-mediated reduction in context-associated seeking behavior is drug and regionally selective. **(A)** HDAC5-3SA in the PrL does not alter sucrose self-administration behaviors nor **(B)** context associated seeking or **(C)** cued reinstatement indicating that HDAC5’s seeking suppression is drug-dependent. **(D)** Experimental timeline. **(E-G)** HDAC5-3SA in the IL had no significant effect on acquisition, context test, or cue test seeking, indicating that the suppressive effect is PrL specific.

The IL is a complimentary mPFC region just ventral to the PrL, and it plays an essential role in extinction of drug-seeking behaviors^17,36,37^. To test whether HDAC5’s ability to limit context-associated cocaine seeking is specific to the PrL, we expressed AAV2-HDAC5-3SA or a control virus targeting the IL region of adult rats (**Fig. 3D**). However, HDAC5-3SA expression in the IL had no significant effects on acquisition of cocaine SA, operant discrimination, total drug infusions, context-associated seeking, extinction learning, or cue-reinstated cocaine seeking (**Fig. 3E-G, S3C-D**). There was a statistical trend for IL HDAC5-3SA to reduced cocaine prime-reinstated drug seeking, but it did not quite reach statistical significance (**Fig. S3E,** t(8)=2.266, *p*=0.0533). Taken together our findings show that HDAC5 functions selectively in the PrL, but not IL, to limit context-associated cocaine seeking.

Since (1) bidirectional PrL HDAC5 manipulations had no effect on cue-reinstated cocaine seeking and (2) the PrL to NAc core (PrL→NAcore) circuit is required for cued reinstatement^15,38–41^, we speculated that HDAC5 does not limit context-associated seeking through suppression of the PrL→NAcore circuit. However, a role of the PrL→NAcore circuit in context-associated cocaine seeking has not been tested. To this end, we used a pathway-specific, inhibitory chemogenetic approach (**Fig. 4A**). Using a Cre-dependent, inhibitory designer receptor exclusively activated by designer drugs (DREADD, DIO-hM4Di-mCherry, or DIO-mCherry control) in the PrL and a retrograde, AAVrg-Cre-GFP in the NAcore, we selectively expressed Gi-DREADD in the PrL→NAcore neurons. Following validation with *ex vivo* acute slice electrophysiology (**Fig. S3A,** t(4)=3.904, *p=*0.0175), we allowed rats to undergo cocaine SA and then administered clozapine N-oxide (CNO; 5 mg/kg; *i.p.*) to suppress PrL→NAcore circuit activity during the context-associated cocaine-seeking test. Suppression of PrL→NAcore activity failed to alter context-associated cocaine seeking (**Fig. 4D**), but in agreement with the prior literature^15,42^, chemogenetic inhibition of the PrL→NAcore pathway in these same rats suppressed cue-reinstated cocaine seeking (**Fig. 4E,** t(26)=3.936*, p=*0.0006), confirming the efficacy of our chemogenetic approach. These findings reveal that the PrL→NAcore is not required for context-associated cocaine seeking and that HDAC5 must limit this mode of cocaine seeking through a PrL→NAcore-independent circuit.

**Figure 4.**
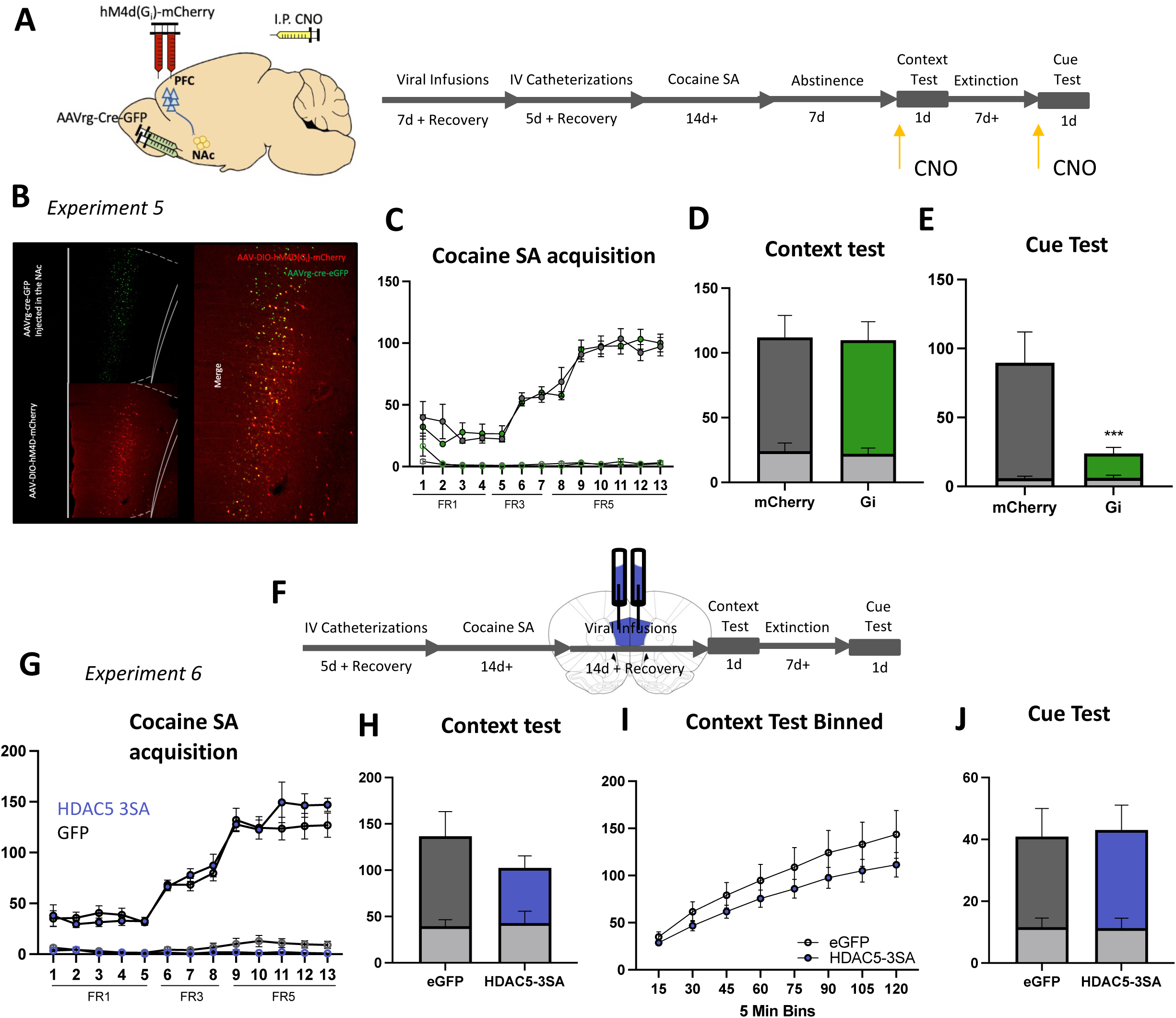
HDAC5-3SA acts selectively in the PrL, during acquisition of drug-taking behavior, to reduce context associated seeking. (**A**) Experiment 5 timeline showing pathway specific chemogenetic viral strategy with hM4d(Gi) expression in NAc-projecting PrL neurons to assay the circuitry underlying context associated seeking. (**B**) Representative expression of GFP-tagged AAVrg-cre in NAc projecting PrL neurons, mCherry tagged hM4D+ neurons, and overlay(**C**) There were no group differences in IVSA behavior. **(D)** Following 7 days of abstinence, CNO-mediated inhibition of NAc-projecting neurons failed to alter context associated seeking behavior but (**E**) did significantly suppress cue-induced reinstatement following extinction to criteria. **(F)** Experiment 6 timeline with cocaine self-administration acquisition prior to viral expression. 24-48 hours following the last day of stable cocaine SA, animals were infused with AAV-HDAC5-3SA and underwent three weeks of forced abstinence to allow for sufficient AAV expression. **(H)** HDAC5-3SA expressed following acquisition did not suppress context associated cocaine seeking, **(I)** with no effects observed across the full 2-hour session (in 5 min bins), and **(J)** no effect on cue reinstatement, indicating that HDAC5-3SA works during drug-acquisition to dampen context-drug associations.

Since PrL HDAC5-3SA suppressed context-associated cocaine seeking at 1 day of withdrawal, we speculated that HDAC5 might be functioning during active cocaine SA to limit formation of drug-context associations. To test this idea, we allowed rats to SA cocaine for 2 weeks prior to expressing HDAC5-3SA bilaterally in the PrL (**Fig. 4F**). We then allowed for 2 weeks of withdrawal to allow for HDAC5-3SA to express prior to testing them in the context-associated cocaine seeking test (i.e., extinction day 1). Prior to virus infusion, there were no significant differences in lever pressing or in cocaine infusions between the assigned groups (**Fig. 4G**, **S3D**). Unlike the expression of PrL HDAC5-3SA throughout acquisition of cocaine SA (**Fig. 2**), we observed no significant effects of PrL HDAC5-3SA on context-associated cocaine seeking when it is expressed after completion of cocaine SA (**Fig. 4H**; t(29)=1.107; *p*=0.2774). To further ensure that we were not missing an effect on drug seeking, we analyzed 5-min bins across the session, but found no significant group differences (**Fig. 4I**). Similar to the PrL HDAC5-3SA study in **Fig. 2**, the expression of PrL HDAC5-3SA after completion of cocaine SA had no effects on extinction learning or cue-induced reinstatement of cocaine seeking (**Fig. S3E and Fig. 4J**). Together these results strongly suggest that PrL HDAC5 limits future context-associated seeking by antagonizing the formation and/or stability of the drug-context association produced during active drug use, but without impacting the reinforcing effects of cocaine or operant discrimination learning and memory.

### HDAC5 influences expression of a select population of cocaine SA-regulated genes

As an epigenetic enzyme, we speculated that PrL HDAC5-3SA influences the formation of drug-context associations through regulation of gene expression. To explore this idea, we expressed HDAC5-3SA or GFP in the PrL and rats were allowed to SA cocaine or saline for 2 weeks followed by 1-week of home-cage abstinence to match the timepoint just prior to the context-associated seeking test. Following bulk RNA-sequencing analysis of PrL tissues, we analyzed differentially expressed genes (DEGs) in these four groups (**Supplemental Table 1)**. EGFP-cocaine SA vs. EGFP-saline SA produced only 44 significant DEGs (**Fig. 5A-B**), and even fewer DEGs (16) produced by HDAC5-saline SA vs. EGFP-saline SA rats. However, we detected the largest number of significant DEGs (113) in the HDAC5-cocaine SA vs. HDAC5-saline SA group, revealing an interesting interaction between cocaine SA and HDAC5 on gene expression. Similarly, HDAC5-cocaine SA produced 64 DEGs compared to EGFP-cocaine, again revealing an interesting interaction between HDAC5 and cocaine SA experience. Moreover, the majority of the HDAC5 DEGs were downregulated, suggesting that cocaine SA experience upregulates a population of genes and HDAC5-3SA represses the expression of a subset. Gene ontology analysis revealed that HDAC5-cocaine had the strongest effect on ‘modulation of chemical synaptic transmission’ in comparison to HDAC5-saline and eGFP-cocaine (**Fig. 5C**) and that under cocaine conditions, HDAC5-3SA reduced expression of glutamatergic signaling pathways genes (**Fig. 5D**), indicating that excitatory/inhibitory balance may be affected. These downregulated DEGs (**Figs. 5F,G**) include important neuronal plasticity genes, such as the NMDA receptor subunit gene, *Grin2b*, and a voltage-gated calcium channel gene, *Cacna1e*. While the importance of these gene for drug-context associations remains unclear, the findings reveal an intriguing interaction between HDAC5 and cocaine SA experience on multiple plasticity genes in PrL that might control the formation and/or stability of drug-context associations produced during active drug use.

**Figure 5.**
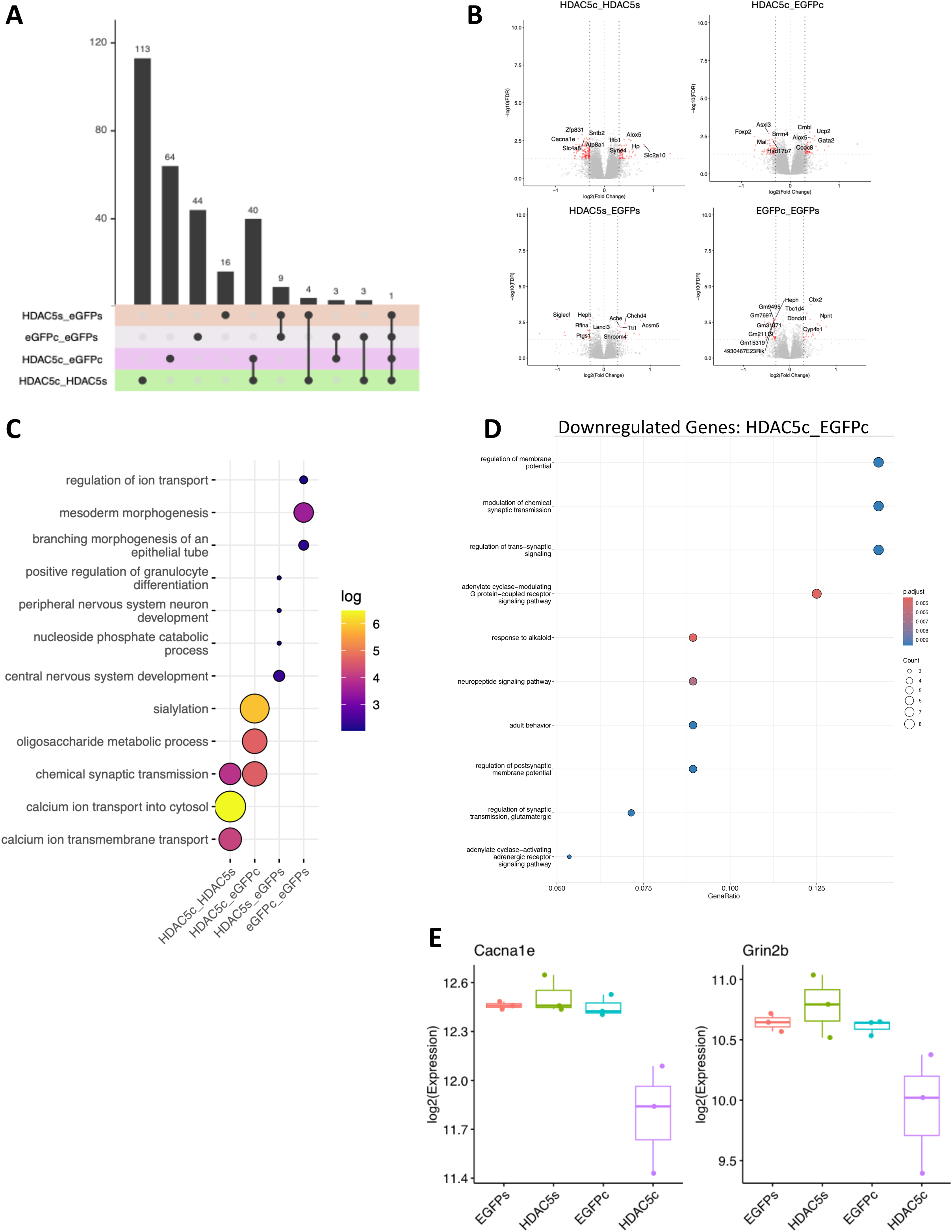
HDAC5 alters cocaine SA-induced gene expression in genes associated with excitatory and inhibitory neurons. (A) UpSet plot depicting overlapping DEGs between cocaine SA vs saline SA groups, +/-HDAC5-3SA. (B) Volcano plots showing the distribution of significant DEGs (FDR<0.05; log2 (FC)>|0.3|). (C) Bubble chart depicting gene ontology analysis by group. (D) Bubble plot showing gene ontology analysis between HDAC5 and eGFP under cocaine conditions. (E) Box plot of expression levels of genes relevant to cocaine seeking and suppressed by HDAC5 following cocaine SA.

### HDAC5 enhances synaptic inhibition and reduces SA context-induced activation

Because HDAC5 influences synaptic signaling genes, we tested the influence of HDAC5-3SA on the synaptic properties of PrL deep-layer pyramidal neurons. We performed acute slice recordings of layer V pyramidal neurons (L5 PNs) expressing HDAC5-3SA or EGFP only. Using patch-clamp configuration, we observed that HDAC5-3SA produced a significant increase in electrically evoked inhibitory postsynaptic currents (eIPSCs) on L5 PNs (**Fig. 6A**, Group main effect, F(1,34)=8.629, *p***=**0.0059), but there was no significant effect of HDAC5-3SA on electrically evoked excitatory postsynaptic currents (eEPSCs) (**Fig. 6B**). Indeed, analysis of the AMPA/GABA ratio on L5 PNs revealed that HDAC5-3SA produced a significant main effect of HDAC5-3SA to reduce the excitatory/inhibitory (E/I) ratio (**Figs. 6C,D**, F(1,34)= 4.875, *p*=0.0241).

**Figure 6.**
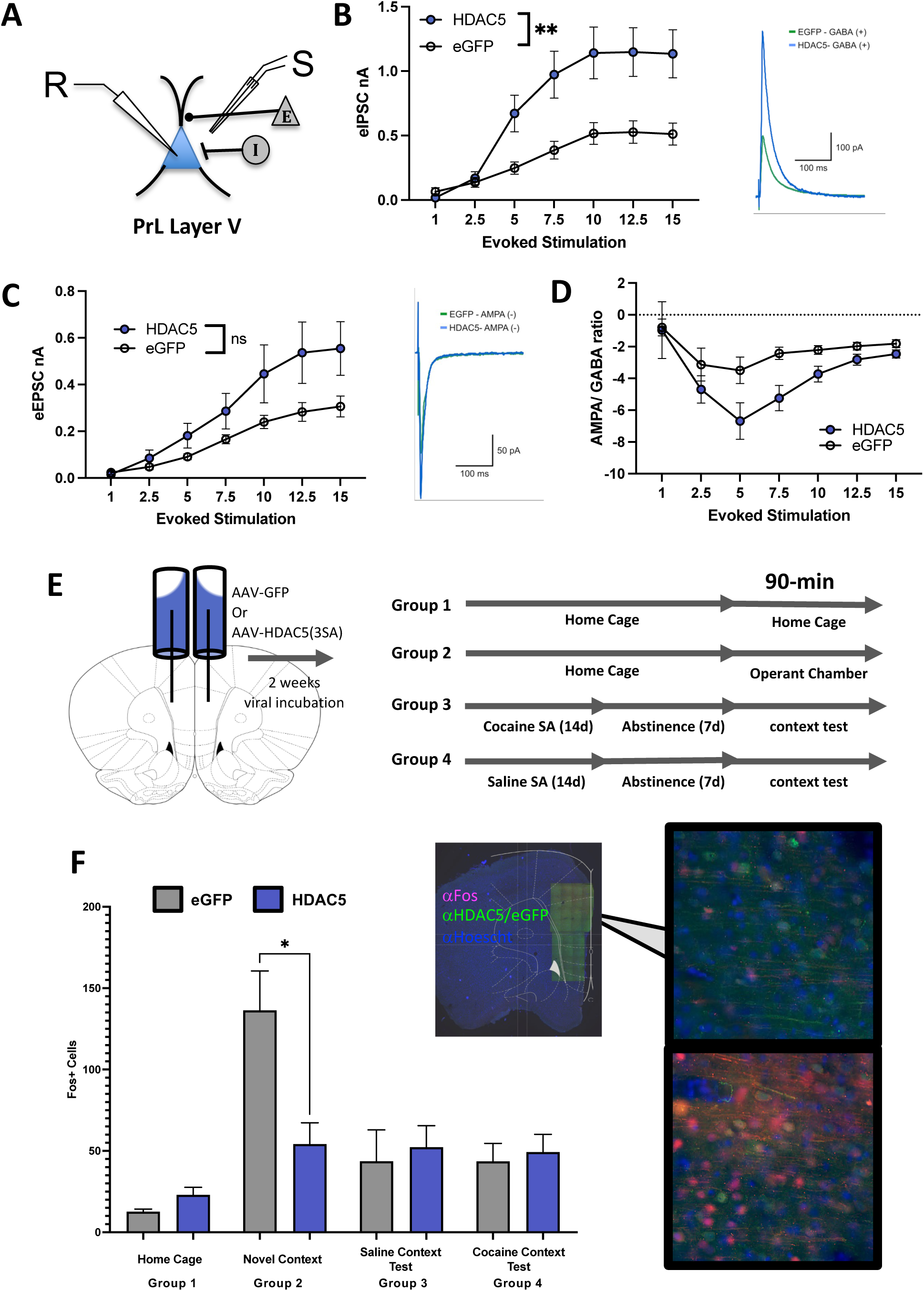
HDAC5 enhances GABAergic transmission on PrL pyramidal neurons and reduces FOS induction during novel environment exposure. (**A**) Experimental design of patch clamp electrophysiology in PrL neurons +/-HDAC5-3SA. **(B)** Cortical HDAC5-3SA expression causes an increase in GABAergic tone in layer V pyramidal neurons, while not altering AMPA related responses. Stepwise stimulation of pyramidal neurons revealed an increased IPSC amplitude, but no significant change in EPSC-evoked amplitude (*p*=0.092). **(C-D)** This resulted in an overall decrease of AMPA/GABA ratio driven primarily by increased GABA evoked stimulation. **(E)** Experimental design for Fos neuronal activation assessment following exposure to a novel context, a context test (following either cocaine or saline SA), or a home cage control. **(F)** HDAC5-3SA inhibited FOS expression only in animals exposed to the Novel Context.

Since a reduction in the L5 PN E/I ratio might impact the ability of the drug-use context to activate the PrL, we measured the effects of HDAC5-3SA on the induction of the immediate early gene, *c-fos*, following introduction to the operant chamber as a proxy for neural activity in this region^43,44^. Rats were infused in the PrL with AAV2-HDAC5-3SA or AAV2-EGFP control and allowed 2 weeks for viral expression. We compared PrL FOS induction in SA-naïve rats taken from their home cage (Group 1; baseline FOS-levels), rats exposed to the SA operant chamber for the first time (Group 2; novel context experience), rats trained to SA cocaine re-exposed to the operant chamber after 1-week of home-cage abstinence (Groups 3; context-associated seeking test), and yoked-saline SA rats re-exposed to the operant chamber after 1-week of home-cage abstinence (Group 4; control group for cocaine SA) (**Fig. 6E**). Interestingly, HDAC5-3SA had no effects on PrL FOS-positive density after cocaine or saline SA in the PrL, but it dramatically reduced the induction of the PrL FOS ensemble in the SA-naïve rats exposed to the operant chamber for the first time (Group 2; **Fig. 6F**, F(7,41) = 5.229, *p=0.0003*, planned multiple comparisons, eGFP novel context vs. HDAC5-3SA novel context, *p=*0.0214). Considering that HDAC5 selectively reduced context-associated seeking only when expressed during active cocaine SA, these findings suggest that PrL HDAC5 limits the strength of neural activity induced by the operant-chamber environment, which might be critical for synaptic plasticity supporting the formation of drug-context associations.

## Discussion

A major obstacle in treating SUD patients is the persistent and prepotent nature of drug-cue and drug-context associations that can trigger a return to active drug use. However, there remains an incomplete understanding of the brain mechanisms by which drug experiences couple to the drug-use environments. In the present study, we examined how the epigenetic enzyme, HDAC5, acts within the mPFC to influence a highly relevant relapse-like behavior: drug-seeking produced by re-exposure to the former drug-use environment. We found that reduction of endogenous HDAC5 in the PrL selectively enhanced context-associated cocaine seeking, but without impacting acquisition of cocaine SA, extinction learning, or cued or drug-primed reinstatement of cocaine seeking. Viral-mediated overexpression of a nuclear-enriched HDAC5 protein, HDAC5-3SA, in PrL, but not IL, had the opposite effect by reducing context-associated cocaine seeking after 1-or 7-days of home-cage abstinence, but again, without altering other aspects of cocaine SA or reinstated cocaine seeking. PrL HDAC5-3SA had no effects on sucrose SA or context-associated sucrose seeking, and it had no effects when manipulated after the establishment of stable cocaine SA, suggesting a selective role for HDAC5 in the formation and/or stabilization of drug-, but not natural reward-, context associations and drug seeking produced by exposure to the drug-use environment. We also showed that, unlike cue-reinstated cocaine seeking, the PrL→NAcore pathway is not required for context-associated cocaine seeking, suggesting that HDAC5 influences a distinct PrL-dependent circuit to influence this mode of relapse-like drug seeking. Analysis of HDAC5-regulated gene expression suggests that it impacted the cocaine SA-regulated gene expression of several ontological gene groups linked to neural plasticity and function. Finally, we found that PrL HDAC5-3SA reduced the E/I ratio of synaptic transmission onto deep-layer PNs, largely by significantly increasing GABAergic inhibitory transmission, and it reduced the novel context-induced activation of PrL, as measured by induction of the FOS-positive ensemble in this region. Taken together, our findings revealed a novel role for HDAC5 in the PrL, but not IL, to limit cocaine, but not sucrose, seeking produced by re-exposure to a former drug-use environment, and that the strength of drug-context associations are encoded, at least in part, by epigenetic mechanisms in the PrL.

Many preclinical drug-seeking studies focus on reinstated drug seeking after extinction of the drug use environment, but our study focused on an understudied mode of relapse-like behavior – cocaine seeking triggered by re-exposure to a former drug-use environment under extinction conditions. There is surprisingly little known about the brain circuits involved in this form of drug seeking. The projections from the PrL→NAcore are well-described as being essential for discrete cue-reinstated drug seeking^15,40,42,45^, but surprisingly, this projection is dispensable for cocaine seeking induced in the drug-use environment under extinction conditions.

We previously showed that HDAC5 functions in the NAc to limit multiple drug-related behaviors, including cocaine CPP and multiple forms of cocaine-or heroin-seeking behaviors, including cue- and drug prime-reinstated seeking. Similar to the findings in the PrL, HDAC5 in the NAc does not alter sucrose SA or any form of sucrose seeking ^28,30^. In striatal neurons, elevation of cAMP signaling by forskolin, activation of D1 dopamine receptors, or cocaine or heroin exposure induces a transient increase in nuclear HDAC5, which is produced in large part by protein phosphatase 2A (PP2A)-dependent dephosphorylation of HDAC5’s conserved serines^28,30^. The dephosphorylation-dependent nuclear accumulation in striatal MSNs is mimicked by the HDAC5-3SA mutant (i.e., S259A/S279A/S498A) ^29^, and our studies here reveal that HDAC5-3SA is nuclear-localized in mPFC neurons and that cocaine SA produces a modest increase in nuclear HDAC5. Since viral-mediated reduction of HDAC5 augments context-associated cocaine seeking and overexpression of HDAC5-3SA reduces it, our new findings show that HDAC5 functions in multiple SUD-related brain regions to limit relapse-like, drug-seeking behavior, thus positioning this epigenetic enzyme as a central regulator of neuroadaptations produced by repeated drug use that influence relapse vulnerability.

Transcriptional dysregulation following cocaine use is well-documented in preclinical studies of the medial PFC^24,46,47^ and in the homologous region (dorsolateral PFC; dlPFC) in postmortem samples from SUD patients^48–51^. Furthermore, human data indicates that cocaine use results in a transient state of increased transcriptional activation followed by a refractory period of downregulation after the drug has been metabolized^50^, and rodent studies report that withdrawal from cocaine results in an upregulation of histone H3K9 acetylation in the mPFC and changes in associated gene expression^49^. In the present study, we assessed the state of gene expression in animals after stable cocaine (or yoked saline) SA and 1 week of home-cage abstinence to identify potential stable changes influenced by HDAC5 and/or cocaine SA. Moreover, this same time-point represents the state of the PrL transcriptome just prior to re-exposure to the cocaine SA-paired environment, and since HDAC5 does not appear to function after cocaine SA to limit drug seeking, dynamic transcriptional changes produced during context-associated seeking seemed less relevant to HDAC5’s function in this context. Interestingly, we only detected 44 DEGs in the comparison of EGFP-saline SA vs. EGFP-cocaine SA, suggesting that cocaine SA plus 1-week of abstinence has a modest impact on PrL gene expression. However, in the HDAC5-3SA-cocaine SA animals, we observed larger effects on PrL gene expression compared to EGFP-cocaine SA (i.e., 64 DEGs) or compared to HDAC5-3SA-saline SA (i.e., 113 genes). Indeed, analysis revealed two intriguing DEGs where HDAC5-3SA reduces the expression only when animals had a history of cocaine SA, including *Grin2b*, which encodes the NMDA receptor subunit implicated in synapse plasticity, learning and memory, and cocaine use disorder, and *Cacna1e*, a voltage-gated R-type calcium channel subunit that gates calcium entry into neurons to influence their firing patterns^52,53^. Future studies will be important to examine the functional relevance of these HDAC5-3SA DEGs, but they suggest a potential means by which HDAC5 might limit PrL plasticity supporting drug-context reactivity and context-associated cocaine seeking. In particular, it will be important to understand how HDAC5-3SA regulated increase in GABAergic synaptic transmission onto PrL layer 5 PNs, and to determine cell autonomous vs. non-autonomous roles for HDAC5 in PrL.

Exposure to the environments in which drug use occurred is oftentimes unavoidable, and the circuitry of diffuse contextual cues underlying drug-seeking behavior is poorly understood. Many cocaine-seeking paradigms focus on reinstated cocaine-seeking after a form of extinction (contextual or cued)^54–56^. In these paradigms, the context-associated seeking test is considered ‘extinction day one’ to adequately resolve cue-induced reinstatement from general SA context-associated seeking. Our findings indicate that this context-associated seeking has unique and separable drivers from cue- or drug prime-reinstated cocaine seeking, as exhibited by HDAC5’s selective regulation of this behavior. Previous studies indicate the regions underlying extinguished contextual cocaine seeking and contextual cocaine seeking after abstinence are distinct. For example, pharmacological inactivation of the dmPFC prevents context-induced cocaine seeking after extinction in an ABA paradigm (whereupon extinction training occurs in a distinct environment to the original cocaine SA acquisition), but not following forced abstinence and re-exposure to the cocaine SA context, as tested in the present study^57,58^. We similarly found that chemogenetic inhibition of the PrL→NAcore pathway did not reduce context-associated seeking. Indeed, our results indicate that the HDAC5-mediated effects in the PrL during the context-associated seeking test were dispensable in comparison to HDAC5’s role during self-administration training. These results are strengthened by our finding that PrL activation, as measured by FOS+ cells, was not different between HDAC5-3SA-expressing rats and controls following context-associated seeking, but was significantly different following first exposure to the IVSA chambers. Further, we found that HDAC5-3SA did not suppress sucrose seeking, indicating that, while HDAC5 inhibited novel context-induced PrL activation, cocaine SA experience was required for HDAC5-mediated long-term effects on seeking, which suggests an interaction between context-induced PrL activity and drug-selective experiences.

Taken together, our findings show that HDAC5 acts within the PrL to enhance inhibitory tone and modulate cocaine-induced gene expression, resulting in dampened context-drug associations and reduction of future context-associated cocaine seeking. These data provide important insight into an understudied modality of drug-seeking: the unextinguished drug-use context. Future studies will focus on understanding the circuitry involved in unextinguished context-associated seeking and determining the important regions upstream and downstream of the PrL in this form of drug-seeking.

## Material & Methods

### Cell Culture and Transfection

(**Fig1A**): Primary embryonic (E18/19) cortical neurons were extracted from Long Evans Rats (Charles River) as previously described^59^ and plated onto slides in a 12 well plate. They were incubated with pre-warmed RM, Neurobasal(NB)/B27, and NB media. Plated cortical neurons were transfected with hHDAC5-eGFP^28^ using CaCl2 transfection as previously described^28^. Briefly, wells were washed with 1x NB. [3.3]ng DNA/[2.5]M CaCl2/ 2x HBSS mixtures were prepared and 50ul were added drop-wise to each well. Wells were incubated for 12 mins at 37°C, 5% CO2, then washed with 0.5mls of recovery medium (BME/ 10%FBS/Penstrep/Q) + 50um APV and incubated for 1 hour in 0.5mls of Recovery medium +50um AP5. Wells were then washed with Rec Med and Neurobasal/B27(PS/Q) and incubated for 42 hours in fresh NB/B27/PS/Q at 37°C.

### Subcellular Localization Analysis

#### In vitro Analysis

(Fig 1A) ∼48hrs after transfection, wells were stimulated 10um of Forskolin or DMSO and incubated for 3 hours. They were then washed with 0.5mls of ice-cold PBS and fixed with 5mls of 4% formaldehyde, 2% sucrose in PB, incubated for 20mins, and washed 3 times with PBS and incubated at 4C for 3 days. Cells were permeabilized and blocked with 0.4% Triton X-100, 10% non-fat dry milk, 1% normal donkey serum in TBS-Tween 0.05% for 1 hour at room temperature and washed twice with PBS. Anti-GFP (chicken, 1:10,000, Aves GFP-1020) or Anti-HDAC5-35 (mouse, 1:200, Abcam ab50001) in 3% BSA TBST (0.05%) was added and incubated overnight at 4C, and washed 3x with PBS. Then wells were incubated with goat anti-chicken 488 (1:500 Abcam ab150169) or anti-mouse 488 (1:500 Abcam ab150113) in 3% BSA TBST (0.05%) overnight at 4C. Hoechst stain (1:10,000) was added in PBS and incubated for 1 min at RT following by 3x PBS washes. Coverslips were mounted on microscope slides and secured with Aquamount and imaged on an epifluorescent microscope (Leica DFC 9000 GT). Transfected cells were identified using mCherry expression, and localization was determined by comparing GFP tag with Hoechst counter stain at 20x power using a Nikon Eclipse 80i fluorescent microscope by a blinded experimenter. Cells were categorized as nuclear or cytoplasmic. A mixed distribution was considered cytoplasmic. Localization is expressed as the percentage of each subcellular localization of the total cells measured.

#### In vivo Analysis

(Fig 1B): Rats expressing 3x-Flag-HDAC5-3SA or WT in the PrL were transcardially perfused with cold 4% paraformaldehyde and 1xPBS flush. 60um sections were taken on a Leica Vibrating Microtome. See below for IHC and antibody details. Images were acquired with a Zeiss LSM 880 confocal microscope equipped with a 63x oil immersion objective (1.4 NA), 0.2um step size, 1.5x digital zoom. Images were then deconvolved with Zen Black (v14, Carl Zeiss) with automatic filtration settings. Images were then imported into Imaris (v9, Bitplane). A surface was built around the nucleus using the DAPI signal. We then expanded that surface by 2.5 um to make a new surface (cytoplasm) using the “Dilate-Surfaces” MatLab extension. We found this expansion amount most accurately encompassed the cytoplasmic area of the cell without including neighboring cells or dendrites. We then created a surface of the nuclear flag signal (total HDAC5 signal in the nucleus) and a surface of the cytoplasmic flag signal (total HDAC5 signal in the cytoplasm) by masking out flag signal outside the selected surface. The intensity sum of each region was divided by the volume of that region and expressed as the proportion of cytoplasmic to nuclear signal. 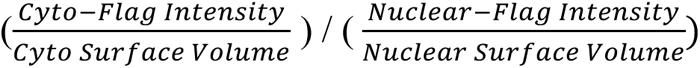.

### Animals

Male Sprague Dawley rats, singly housed upon arrival, aged 6-8 weeks and weighing 275-325g at delivery were kept in a climate-controlled vivarium on a reverse light dark cycle to allow for testing during their active period. Prior to experimental procedures, animals were habituated for a minimum of 3 days and allowed *ad libitum* access to standard chow and water. Animals were 7-9 weeks old at the time of first surgery. During the contingent self-administration portion of the experiments, animals were restricted to 20g of chow per day, or approximately 85% of their free feeding weight. All experimental procedures were approved by the Institutional Animal Care and Use Committee at Medical University of South Carolina, and in accordance with NIH and National Research Council guidelines.

### Viruses

Endogenous HDAC5 expression in the PrL was knocked down with AAV2-shHDAC5-GFP: shHDAC5 Sequence: GAAGGTTCTACAGAGAGCGAG under a U6 promoter, GFP under cytomegalovirus (CMV) promoter produced by the University of South Carolina Vector Core (titer 1.2 X 10^12^, previously published^30,60^), compared to control AAV2-shLuc-GFP: Sequence: CTTACGCTGAGTACTTCGA under a U6 promoter, GFP under cytomegalovirus (CMV) promoter produced by the University of South Carolina Vector Core (titer 1.2 X 10^12^). We used the same *AAV2-HDAC5-3SA:* Promoter: CMV, as previously described ^28^, with an added N-terminal Flag-tag for optimized IHC (titer 6.77x10^14^), and compared to AAV2-GFP(titer 6.07X10^14^), both produced by the University of South Carolina Vector Core. Chemogenetics were conducted using either AAV5-hSyn-DIO-hm4d-mCherry (AddGene #44362-AAV5, titer 1.4x10^13^), AAV5-hsyn-DIO-mCherry (AddGene, #50459-AAV5, titer 1.6x10^13^) in the PrL and AAVrg-EGFP-cre (AddGene #105540-AAVrg, titer 1.17x10^13^).

#### Drugs

<u>CNO *N*-Oxide</u> (Tocris, #4936, reconstituted with 0.9% (w/v) normal saline to 5.0 mg/ml, light protected until i.p. injections).

<u>Cocaine-HCL</u> (NIDA Drug Supply Program) was dissolved in sterile 0.9% (w/v) saline to a final concentration of 4mg/ml to be administered at a dose of 0.5mg/kg/infusion in the rat IVSA paradigm.

### Surgical Procedures

Intracranial Infusions: Viral infusions were delivered stereotaxically as previously described^61^. Rats were anesthetized using 5% Fluriso^TM^, isoflurane (Vet One), and maintained under deep anesthesia at 2-3%. Supportive heat and fluids were provided. Ketoprofen (5mg/kg, s.q.) was administered pre-operatively and cefazolin (10mg/kg s.q.) were administered post-operatively (Patterson Veterinary Supply, Devens MA). Viruses were infused bilaterally into the PrL (+2.8 A/P, +/-0.62mm M/L, -3.8mm D/V), IL (+2.8 A/P, +/-0.64mm M/L, -5.5mm D/V or NAc Core (10° angle; from Bregma: +1.6 A/P, +/-2.8 M/L, -7.1 D/V) with Nanoject Auto-nanoliter Injectors (Drummond Scientific Company, Broomall, PA), and pipettes were left in place for 5 minutes to allow diffusion of virus.

Catheter Implantation: Catheter implantations were performed as described previously^61^. In brief, on end of a silastic catheter (Thermo Fisher Scientific or SAI Infusions Technologies, Lake Villa, IL) is inserted into the right jugular vein, secured, and threaded subcutaneously to a back mounted infusion cannula (Plastics One, Roanoke VA) or Catheter Access Buttons^TM^ with Cannulock^TM^ (SAI Infusions Technologies, Lake Villa, IL).

### Cocaine IVSA

As previously described^61^, rats were trained to self-administer cocaine intravenously in operant chambers (Med-Associates) on an escalating FR schedule (FR1, 3, 5) for 2-hr sessions for approximately 12-14 days. Rats were trained to press an active lever to receive an infusion of cocaine HCL (0.5mg/kg/infusion) paired with a light-tone cue complex. Following a reinforced active lever press, a 15 second time out occurred where presses were recorded but not rewarded with drug infusion. Inactive lever presses were recorded but not reinforced. Self-administration criteria was set at a minimum of 10 cocaine infusions for five days (FR1), 3 days (FR3), and 5 days (FR5). Following completion of FR training, rats were subjected to a period of abstinence in the vivarium (24hrs or 7 days). Following abstinence, rats were returned to the chambers under extinction conditions. Lever presses were recorded but not reinforced with drug or cue complex presentation. When rats reached the extinction criteria of a minimum of 7 days of extinction training and less than 25 active lever presses for 2 consecutive days, they underwent a cue-induced reinstatement test, whereupon pressing of the active lever resulted in the presentation of the cue+tone complex but not a drug infusion. In some experiments, rats were then extinguished to criteria and given a cocaine prime (10 mg/kg, i.p.) prior to being placed in the SA chamber and responses were recorded under extinction conditions.

### General Immunohistochemistry

Following cessation of testing, rats were anesthetized and transcardially perfused with 150 mls of ice-cold phosphate buffered saline (1X PBS) followed by 4% (w/v) PFA (in PBS) at a rate of 20 ml/min. Brains were then post-fixed for 24-hrs in 4% (w/v) PFA (in PBS) and transferred to 30% (w/v) sucrose (in PBS) for cryoprotection, and then sectioned (40um) on a freezing microtome. Sections were then blocked with 3% NDS or NGS (Jackson Imm, 0.3% (v/v) Triton X-100, 0.2% (v/v) Tween-20, 3% (w/v) BSA for 3 hours at RT, and then incubated O/N at 4°C with blocking solution and primary anti-c-Fos (226 008, Synaptic Systems, rabbit, 1:500), anti-mCherry (NBP2-25157 Novus Biologicals, rabbit, 1:1000), anti-Flag (50-165-7385 Sigma Aldrich, mouse, 1:1000), anti-GFP (Aves, GFP-1020, GRP-1026, chicken, 1:10,000, serial diluted), anti-HDAC5 (2082BV, Cell Signaling, Rabbit 1:200, Figure 1 only), anti-HDAC5-35 (Mouse, ab50001 abcam, Mouse (1:500-1:1000) and gentle agitation. Sectioned were then washed 3x 5mins in 1x PBS and incubated for 1 hour under light protected with Alex Fluor^®^ 488 donkey anti-chicken (703-545-155, 1:500, Jackson ImmunoResearch), Alex Fluor^®^ 488 donkey anti-mouse (715-586-150, 1:500, Jackson ImmunoResearch), Alex Fluor^®^ 594 donkey anti rabbit (711-587-003, 1:500, Jackson ImmunoResearch). Sections were washed 3x 5mins in 1xPBS, then incubated with Hoechst 33342 (H3570, 1:10,000 (serially diluted), Invitrogen) for 1 min, washed twice more, then mounted on SuperFrost+ with ProGold Antifade or FluoroGold mount.

### Electrophysiological Procedures

Acute coronal slices (300-µm thickness) containing the PrL cortex were obtained using a Leica VT1200s slicer in semi-frozen artificial cerebrospinal fluid (ACSF) containing (in mM): 127 NaCl, 2.5 KCl, 1.2 NaH2PO4, 24 NaHCO3, 11 D-glucose, 1.2 MgCl2, and 2.40 CaCl2, 0.4 Na-ascorbate (pH 7.4, 315-320 mOsm). Kynurenic acid (5 mM) was added to the ACSF to avoid overactivity of glutamatergic receptors during recording. Slices were then transferred to normal ACSF (without kynurenic acid) to recover at 37°C for 30 minutes, and then transferred to room temperature ACSF for an additional 30-minute recovery, followed by recordings. All solutions were continually equilibrated with 95% O2 and 5% CO2. Pyramidal neurons in Layer V (LV) of PrL were tested and visualized with infrared differential interference contrast optics (DIC/infrared optics) and identified by their location and shape of the neurons and their apical dendrites. Unless stated otherwise, all electrophysiological experiments were performed in whole cell voltage clamp mode at -70 mV using borosilicate pipettes (4-6 MΩ) manufactured by Narishige puller or pulled using NARISHIGE puller (NARISHIGE, PG10) from borosilicate tubing (Sutter Instruments) and filled by an internal solution containing (in mM): 140 CsMeS (CH3CsO3S), 5 KCl, 1 MgCl2, 0.2 EGTA, 11 HEPES, 2 NaATP, 0.2 Na2GTP pH 7.2–7.4 (pH 7.2, 290–295 mOsm).

The evoked postsynaptic inhibitory/excitatory responses of neurons in LV of PrL were elicited by field stimulation of excitatory and inhibitory afferents in LV. The low intensity pulses of stimulated current (25–100 μA, 100 μs duration) were applied through a fine-tipped (∼2-3 μm), bipolar stimulating electrode made from borosilicate theta glass capillary tubing (Warner Instruments).

AMPA-receptor-mediated excitatory postsynaptic currents (EPSCs) were recorded in voltage clamp mode at -70 mV (reversal potential for GABA current) to minimize GABAA current. Inhibitory postsynaptic currents (IPSCs) mediated by GABAA receptors were recorded in voltage clamp mode at 0 mV (reversal potential for AMPA currents) to eliminate the current through AMPA-receptors. To calculate E/I (AMPA/GABA) ratio the responses, evoked by field stimulation of bipolar stimulating electrode, the amplitude of the AMPA response recorded at -70 mV was divided by the amplitude of the GABA response at 0 mV.

All recording data were acquired and analyzed by amplifier AXOPATCH 200B (Axon Instruments), digitizer BNC2090 (X National instruments) and software AxoGraph v.1.7.0, Clampfit v.8.0 (pClamp, Molecular devices) and MiniAnalysis Program v.6.0.9 (Synaptosoft). Data were filtered at 2 kHz by AXOPATCH 200B amplifier (Axon Instruments) and digitized at 10-20 kHz via AxoGraph v.1.7.0. software.

### RNA-sequencing

Tissue punches containing the rat PrL were extracted from virally infected tissue using a 2mm biopsy punch. Tissue was processed for RNA extraction using a modified manufacturer’s procedure (miRNA Easy Kit, Qiagen). Briefly, RNA is extracted using phenol/chloroform, bound to the column, and washed with included buffers. During the wash steps, samples were treated for 15 mins at room temperature with RNase-free DNAse1 (Qiagen), followed by final washing and elution. RNA concentration and OD260/280 were determined by nanodrop. Frozen samples were sent to BGI Americas Cooperation (Cambridge, MA) and sequencing was performed using polyA mRNA directional RNA-seq library preparation using a DNBSEQ^TM^ platform.

### RNA-seq alignment and quality control

Quality of raw sequence data was assessed using FastQC (v0.11.9). Reads were aligned to the rat mRatBn7.2 reference genome using STAR (v2.7.1a)^62^. Secondary alignment and multi-mapped reads were further removed using in-house bash scripts. Only uniquely mapped reads were retained for further analyses. Quality control metrics were assessed by Picard tool (http://broadinstitute.github.io/picard/). Ensembl gene annotation for mRatBn7.2 was used as reference alignment and downstream gene quantification. Gene level expression was calculated using featureCounts (v2.0.1)^63^. Counts were calculated based on protein-coding genes from the annotation file.

### Differential expression

Counts were normalized using counts per million reads (CPM). Genes with no reads were removed. To infer potential experimental confounders, we calculated surrogates variables were calculated using the sva package in R^64^. Differential expression analysis was performed using a linear model post-hoc analysis in R using emmeans (v1.34)^65^ with the following model: gene expression ∼ Genotype * Condition. We estimated log2 fold changes and P-values. P-values were adjusted for multiple comparisons using TukeyHSD. Differentially expressed genes where consider for FDR<0.05 and a fold-change ≥ |0.3| as statistically significant. Rat Gene ID were translated into mouse and human Gene ID using biomaRt package in R^66^.

### Gene Ontology (GO) analysis

The functional annotation of differentially expressed and co-expressed genes was performed using clusterProfiler (v4.2) and enrichR (v3.2). A Benjamini-Hochberg FDR (FDR<0.05) was applied as a multiple comparison adjustment.

### Code availability

Codes to support the DGE analysis: https://github.com/BertoLabMUSC/Barry_HDAC5.

### Availability of data and material

The NCBI GEO accession number for the data reported in this manuscript is [in process].

## Data Availability Statement

Data available on request from authors.

## Ethics Approval Statement

All procedures were approved by the Medical University of South Carolina’s Institutional Animal Care & Use Committee. Experiments and analysis were performed following NIH rigor and reproducibility guidelines.

