## Supplemental Figures for "Histone deacetylase 5 in prelimbic prefrontal cortex limits context-associated cocaine seeking"

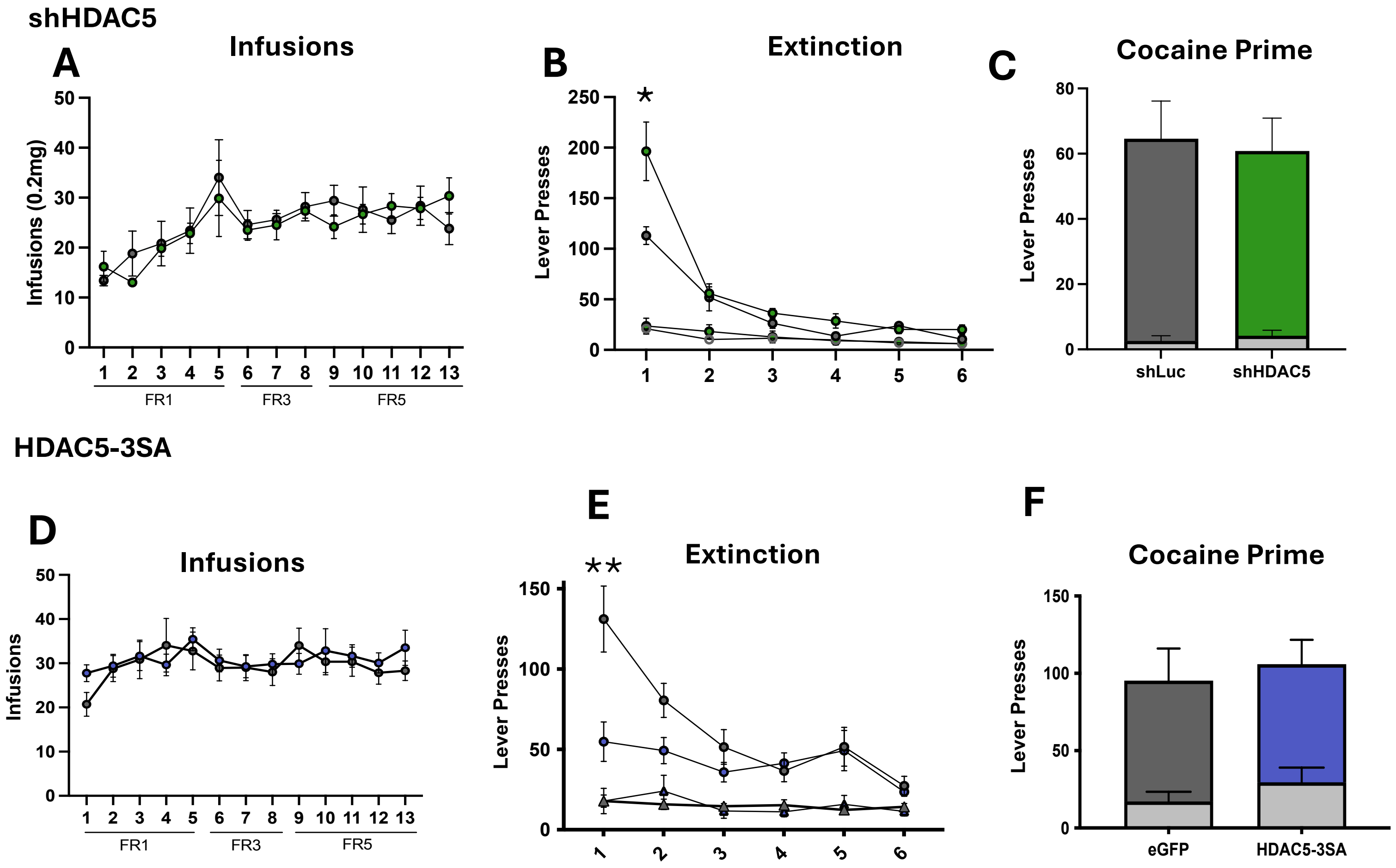


**Supplemental Figure S1.** shHDAC5 expression did not alter total cocaine intake during acquisition (**A**) or alter extinction behavior (**B**) or cocaine-primed reinstatement (**C**). HDAC5-3SA expression in the PrL did not alter infusions, extent of extinction, or cocaine-primed reinstatement (**D-F**).

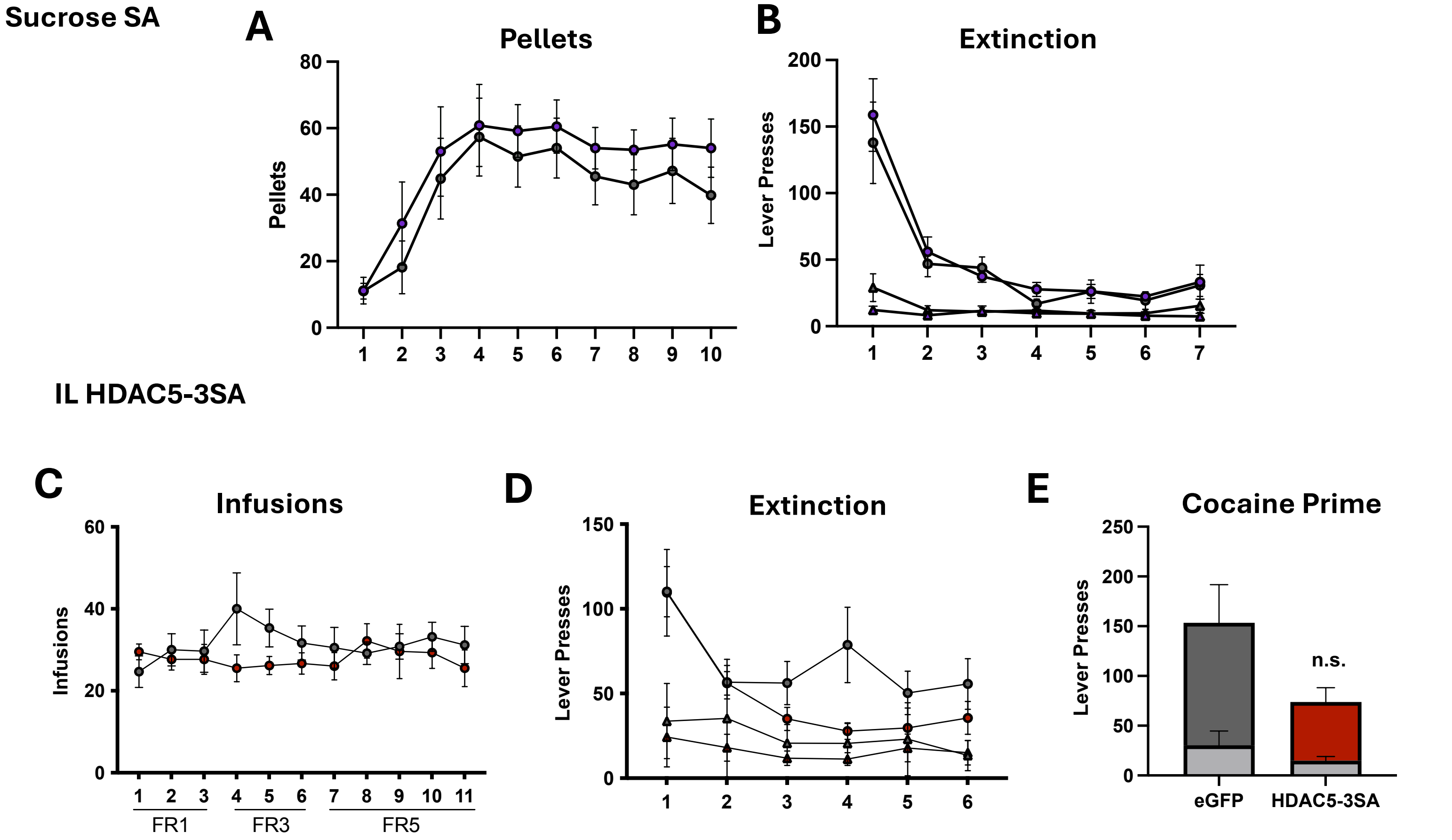


**Supplemental Figure S2.** HDAC5-3SA expression did not alter total sucrose pellets administered in sucrose SA (**A**) or lever-pressing during extinction training (**B**). HDAC5-3SA expression in the IL did not alter total cocaine infusions or extinction behavior (**C-D**). There was a statistical trend for a decrease in cocaine-primed seeking behavior, but these data failed to reach significance (**E**).

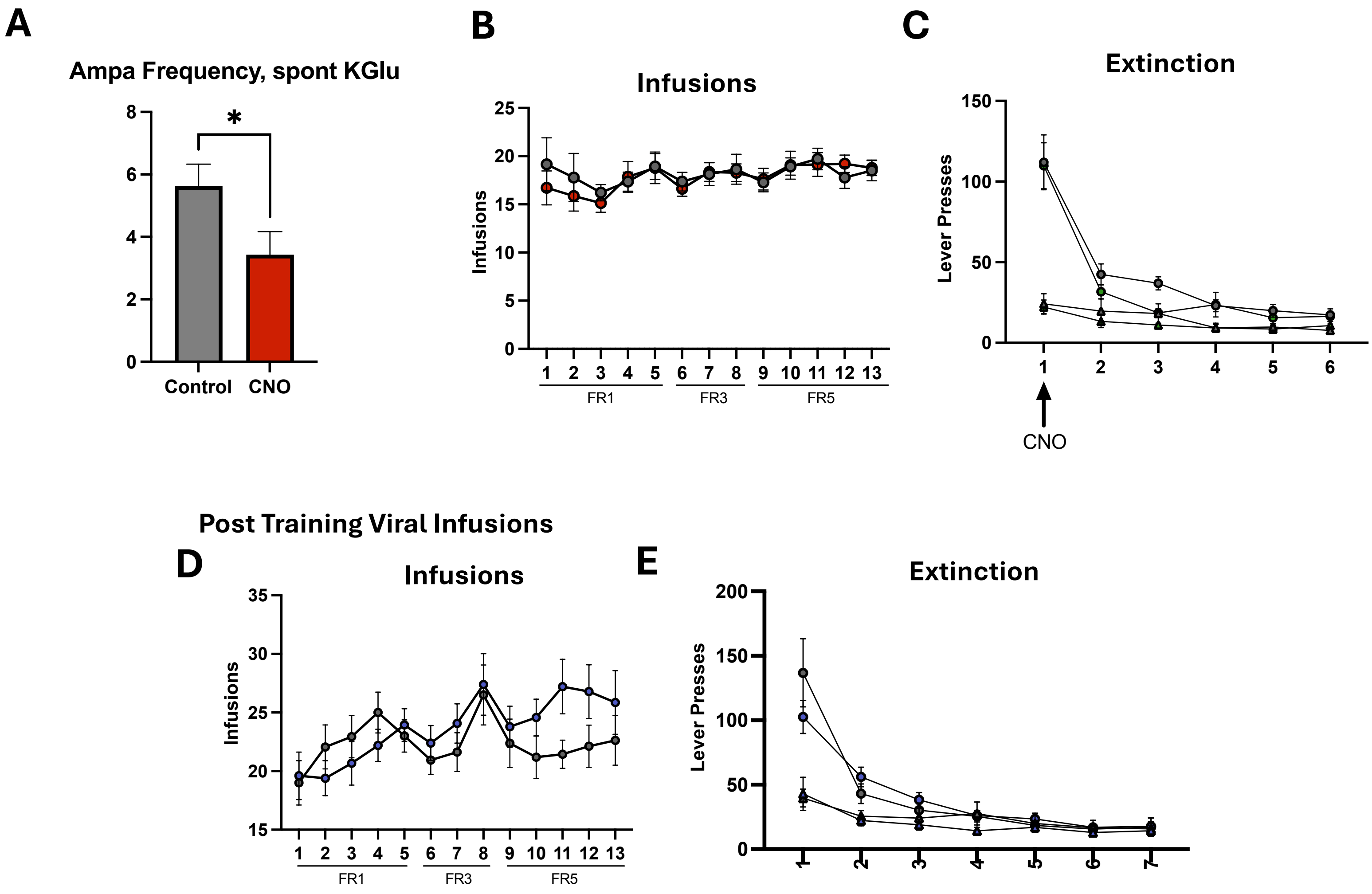


**Supplemental Figure S3.** Acute slice electrophysiological validation of hM4d(Gi) DREADD efficacy in PrL pyramidal neurons showed a reduction in spontaneous AMPA frequency (**A**). Viral mediated DREADD and control expression did not alter overall cocaine intake or extinction behavior (**B-C**). Prior to infusion with HDAC5-3SA, there were no differences in cocaine infusions (**D**) or subsequent extinction behavior (**E**).
